# T helper cells exhibit a dynamic and reversible 3’UTR landscape

**DOI:** 10.1101/2023.01.19.523753

**Authors:** Denis Seyres, Oliver Gorka, Ralf Schmidt, Romina Marone, Mihaela Zavolan, Lukas T. Jeker

**Author notes:** Authors contributed equally (Denis Seyres, Oliver Gorka, Ralf Schmidt).

## Abstract

3’ untranslated regions (3’UTRs) are critical elements of messenger RNAs, as they contain binding sites for RNA-binding proteins (RBP) and microRNAs that affect various aspects of the RNA life cycle including transcript stability and cellular localisation. In response to T cell receptor activation, T cells undergo massive expansion during the effector phase of the immune response and dynamically modify their 3’UTRs. Whether this serves to directly regulate the abundance of specific mRNAs or is a secondary effect of proliferation remains unclear. To study 3’UTR dynamics in T helper cells we investigated division-dependent alternative polyadenylation (APA). We generated 3’ end UTR sequencing data from naïve, activated, memory and regulatory CD4+ T cells. 3’UTR length changes were estimated using a non-negative matrix factorization approach and were compared with those inferred from long-read PacBio sequencing. We found that APA events were transient and reverted after effector phase expansion. Using an orthogonal bulk RNAseq dataset, we did not find evidence of APA association with differential gene expression or transcript usage, indicating that APA has only a marginal effect on transcript abundance. 3’UTR sequence analysis revealed conserved binding sites for T cell-relevant microRNAs and RBPs in the alternative 3’UTRs. These results indicate that polyA site usage could play an important role in the control of cell fate decisions and homeostasis.

## Introduction

As part of the adaptive immune system, naïve T helper cells emerge from the thymus to circulate through the body in search of their cognate antigen presented by a specialised antigen-presenting cell. During this phase, CD4+ T cells are in a quiescent state, mainly dependent on signals conveyed by a T cell receptor (TCR):Major-histocompatibility complex (MHC)-class II interaction and IL-7 (Surh and Sprent, 2008). Upon encountering their cognate antigen and costimulatory signals, T cells start to proliferate and, in the presence of cytokines, can differentiate into effector T cell subsets with specialised roles (Ruterbusch *et al*., 2020; Dong, 2021). Some T cell subsets promote inflammatory conditions, while regulatory T cells (Tregs) have a dampening, immunomodulatory role (Okeke and Uzonna, 2019); (Sakaguchi *et al*., 2020). Both the expansion and the differentiation phases are tightly regulated, on multiple molecular levels. While the transcriptional control has been widely investigated, post-transcriptional regulation is less well understood. By binding to the 3’ untranslated regions (3’UTRs) of key target transcripts, thereby affecting their stability and protein expression, microRNAs (miRNAs) and RNA-binding proteins (RBPs) regulate many cellular processes including the cell cycle, metabolism, and cell differentiation (Fu and Blackshear, 2017; Bartel, 2018; Rodríguez-Galán, Fernández-Messina and Sánchez-Madrid, 2018). Global shortening of 3’UTRs has been observed in rapidly dividing cells, such as cancer cells and activated lymphocytes, suggesting a link between alternative polyadenylation (APA) and oncogenesis and lymphocyte differentiation (Sandberg *et al*., 2008; Mayr and Bartel, 2009). Conversely, differentiation of embryonic stem cells is associated with the opposite trend of general 3’UTR lengthening (Gruber and Zavolan, 2019; Mitschka and Mayr, 2022)(Ji et al., 2009).

Previous work on human and mouse T cells has shown that activation is accompanied by a genome-wide trend toward increased usage of coding region-proximal polyadenylation sites, leading to shorter 3’UTRs (Sandberg *et al*., 2008; Gruber *et al*., 2016). 3’UTR remodelling did not lead to large changes in mRNA and protein levels (Gruber *et al*., 2016). However, subsequent studies revealed that the 3’UTR isoform-specific RBP interactome can affect other processes such as the localization of the protein translated from the mRNA. One example is provided by CD47 isoforms, in which the 3’UTR acts as a scaffold for the binding of the RBP HuR, which facilitates the localization of CD47 to the plasma membrane (Berkovits and Mayr, 2015). In this project, we set out to determine when the dynamic remodelling of 3’ UTRs takes place in the helper T cell life cycle and what implication this process could have for T cell function. We generated 3’UTR end sequencing data and performed pairwise comparisons of functionally annotated APA genes in naïve, activated and differentiated T cells. While 3’UTR shortening events were found to be more frequent upon T cell activation, we identified a substantial number of lengthening events. To further validate the APA sites inferred by 3’ end sequencing, we also generated PacBio long-read sequencing data. While transcript coverage by PacBio reads was sparse, we found a meaningful overlap between the isoforms identified with this technology and by 3’ end sequencing. Next, we sought to determine whether the APA changes are reversible following T cell activation. We identified 1010 genes associated with transient 3’UTR length changes in T helper cells during acute activation and effector phase expansion, meaning that the cells eventually return to a naïve state of polyadenylation site usage when re-entering a resting state as memory T cells. Using an orthogonal mRNA dataset, we observed a limited effect of APA on gene expression and regulation and identified several miRNA families and RNA binding proteins that may act as cis-regulators. Our results extend previous observations of the dynamics of alternative polyadenylation in primary murine T cells and suggest the presence of additional levels of plasticity in the processes controlling 3’UTR alterations allowing the return to a steady-state level of global transcript length.

## Results

### Analysis of division-dependent 3’UTR length changes in murine CD4^+^ T cells

Cancer cell lines and proliferating primary cells have been reported to exhibit alternative polyadenylation events with global 3’UTR shortening (Sandberg *et al*., 2008; Gruber *et al*., 2016). To further investigate the relationship between proliferation, differentiation and APA, we determined the changes of the 3’UTR landscape during the response of murine primary naïve CD4^+^ T cells to stimulation. We purified naïve CD4^+^CD62L^-^ T cells and labelled the cells with a fluorescent dye (Cell Tracer Violet, CTV), which allows the tracking of the number of divisions as the fluorescence intensity becomes 2-fold smaller with each division. Cells were stimulated through anti-CD3/anti-CD28 antibody crosslinking of the T cell receptor *in vitro* leading to activation and subsequent proliferation (**Figure 1A**). After 48 hours, the activated T cells were sorted to high purity (>98%) according to the dye intensity peaks that resulted in three distinct populations: T cells that did not divide yet (peak 0), cells that have divided once (peak 1), and cells that have divided twice (peak 2) (**Figure 1B**). All sorted populations were used for RNA extraction (**Figure 1A, B**). A fourth population consisted in a CTV-labelled naïve CD4^+^ T population (unstimulated) which in parallel was subjected to Trizol-based RNA extraction at the initiation of the proliferation experiment. We analysed the expression of the activation markers CD44, CD62L, CD69 and CD25 on all sorted populations of divided T cells. Each division was characterised by a distinct expression of these markers. Cells in peak 0 expressed the lowest levels of CD44 and displayed heterogeneous expression of CD69 and the high affinity IL-2 receptor CD25. Cells in peak 1 uniformly expressed high levels of the early activation markers CD69 and CD25. Cells in peak 2 showed the highest expression of CD25 and CD44, whereas CD69 expression was lower than in peak 1. Thus, T cells within peaks 1 and 2 underwent full activation, whereas T cells within peak 0 consisted of a more heterogeneous population. Most cells had upregulated CD69 and CD25 but the activation did not result in cell division (yet) whereas other cells had not induced CD69 expression (**Figures S1A, S1B**). To monitor changes in the global 3’UTR landscape of naïve, activated and proliferating T cells with a division-dependent resolution, we prepared 3’ end sequencing libraries from all four populations using the previously established A-seq2 protocol (Gruber *et al*., 2016; Martin *et al*., 2017). To identify A-seq2 read enriched regions, or “peaks”, genome-wide, we used a density-estimation approach (Boyle *et al*., 2008). We retained a peak if it was present in both technical replicates and in at least one biological sample. These peaks were then filtered with polyA sites available in the PolyASite database (Herrmann *et al*., 2019). In total, we retained 43311 peaks, distributed on 10906 genes. For 38.1% of the genes (4159) we identified one site (**Figure 1C**), thus these genes were not experiencing alternative 3’UTR polyadenylation in our model. Nearly half of the A-seq2 peaks (50.4%) were located in 3’UTRs, the remaining being distributed across introns, exons, last exons, 5’UTR and non-coding regions (20.9%, 8.3%, 2.8%, 0.8% and 16.8% respectively). A principal component analysis (PCA) using the log transformed RPM (Reads per million mapped reads) of reads located on A-seq2 peaks showed that a high proportion of the variance was explained by activation, along the first principal component (PC1) (**Figure 1D**). Focusing on activated samples, we observed higher variance between no division (peak 0) T cell samples and one and two-divisions samples (peaks 1 and 2 respectively) than between samples from 1 or 2 divisions (**Figure S1C**). In addition, with the goal to generate a qualitative catalogue of transcripts effectively present, we used a low coverage long-read PacBio sequencing from naïve and 48h activated T cells (no division and 2 divisions). In line with what was observed with the A-seq2 protocol, more than half of the covered genes presented one isoform (**Figure S1D**). 4194, 6152 and 5340 3’ ends (in the naïve, 0 and two divisions samples respectively) colocalized with 3’ ends of transcripts identified by long read sequencing, whereas 6688, 7050 and 6244 were only identified with long read sequencing. Conversely, 14221, 20246 and 20914 3’UTR ends were only identified by A-seq2 (**Figure 1E**). Due to limited coverage, while these data showed highly promising value, they did not cover the whole transcriptome and were only used as an additional informative layer, in particular for visualisation (**Table S1**). We then compared our 3’ end sequencing data to a similar study performed by Hsin and colleagues (Hsin *et al*., 2018). They analysed the effect of miR-155-deficiency in different murine cellular contexts, including CD4^+^ T cells, through differential iCLIP, RNA-seq, and 3’ untranslated region (UTR)-usage analysis with poly(A)-seq. The authors compared naïve CD4^+^ T cells and in vitro activated CD4^+^ T cells stimulated for 24h and 48h. Although the experimental setups were not identical -Hsin and colleagues did not sort 48h activated cells according to cell divisions and the poly(A)-seq protocol they used differs slightly from the A-seq2 protocol -the covered genes and genomic distribution was comparable (Hsin *et al*., 2018) (**Figure S1E**). Similar important activation effects, when comparing naïve to 48h activated cells, have also been observed by analysing the transcriptome (Dölz *et al*., 2022). Finally, we observed an equivalent distribution of the variance when re-analysing the poly(A)-seq data generated by Hsin and colleagues (**Figure S1F**). We illustrate the data we obtained for a specific gene, Psen1 (**Figure 1F**). This example shows how PacBio long read sequencing can identify novel transcripts, which are not in the reference annotation (Gencode v25), and how well the 3’ end of the long read colocalized with 3’ end revealed by A-seq2. Using our newly generated 3’ end sequencing data we observed a clear effect of CD4^+^ T cell activation on 3’ end coverage. We also observed differences between 48h activated no division cells and 48h activated cells after one and two divisions. Identification of more than one putative 3’ end in approximately 60% of genes with A-seq2 signal showed the diversity of transcript abundances during activation and proliferation phases.

**Figure 1.**
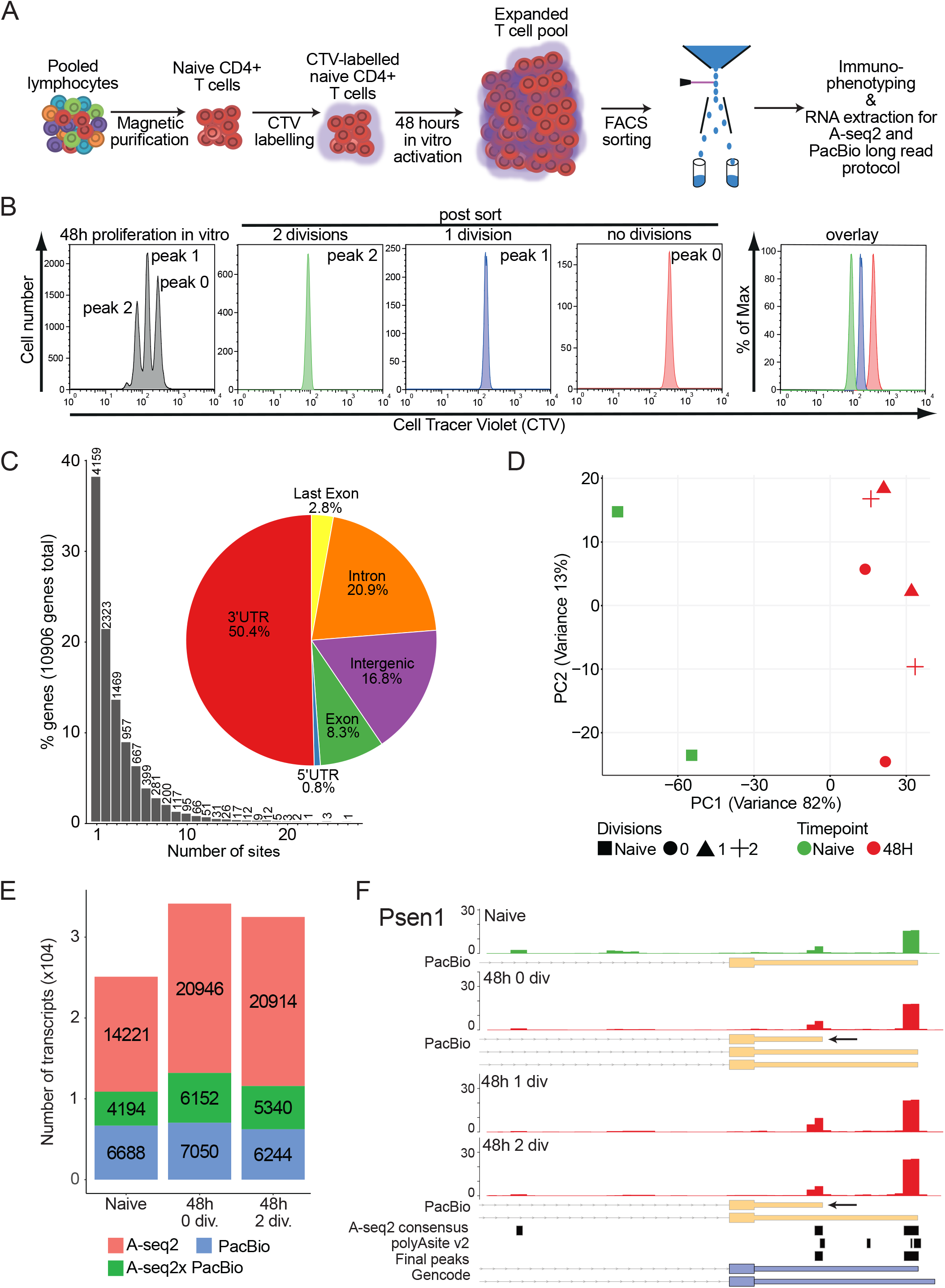
Analysis of division-dependent global 3’UTR changes in murine CD4+ T cells. A) Schematic work-flow of T cell-labeling, activation and sorting. B) Representative CTV-labeling profile of CD4+ T cells after in vitro activation for 48 hours and typical results for sorted peaks of undivided T cells (peak 0), and T cells which are divided once (peak 1) and twice (peak 2). The right panel shows an overlay of all populations after FACS sorting. C) Barplot representing distribution of alternative polyA (APA) sites per genes. Pie plot shows genomic localization distribution of APA sites. D) Principal component analysis (PCA) resulting from read coverage over the final peak set. E) Stacked barplot summarizing 3’UTRs identified by A-seq2 only (red), PacBio only (Blue) or both A-seq2 and PacBio (green). F) Example of Psen1 gene 3’UTR region. Overlap of A-seq2 peaks («consensus») and polyAsite database is labeled as the “Final peaks” track. A-seq2 signal of the different sorted cell populations is represented. Yellow boxes represent isoforms identified using PacBio long read sequencing (arrows indicate newly identified isoforms), blue boxes represent Gencode (v25) isoforms and black boxes indicate PAS.

### Identification of alternative polyA site usage

Changes in 3’UTR length are typically inferred from significant differences in 3’ end read coverage between different putative polyA sites. We decided to use an approach which, at gene-level, uses vector projection and non-negative matrix factorization and allows the identification of changes in tandem polyA sites but also when more alternative polyA sites are located in between most distal and most proximal sites (Yalamanchili *et al*., 2020). For this analysis, we did not consider intergenic sites but, in addition to UTR sites, we included intronic and exonic polyA sites. We devised a pairwise comparison of naïve and all activated samples (**Figure 2A**). Event types were stringently defined to fulfil two conditions: shortening event was defined when sample B proximal projection was greater than sample A proximal projection and if sample B distal projection was smaller than sample A distal projection. The opposite conditions defined a lengthening event. While we observed, on average, a higher number of shortening events, in line with work reported previously about CD4^+^ T cell activation and proliferation (Gruber *et al*., 2014, 2016), we also observed a substantial number of transcripts using more distal APA sites. In addition, a higher number of events were detected when comparing naïve to one and two divisions 48h activated stages (559 and 510 shortening and 437 and 320 lengthening events) than when naïve and activated without division stages were compared (331 and 185 for shortening and lengthening events, respectively). Most of the observed changes were due to APA at tandem poly(A) sites in the same 3’UTR, followed by events involving intronic and then exonic sites (**Figure 2B**). As an example, the short *Ddx46* isoform, whose gene encodes the DEAD-Box Helicase 46 became more prominent upon activation (**Figure 2C**). One of the most proximal sites in this gene accumulated significantly more reads in samples from activated cells, suggesting a higher abundance of the shorter transcript isoform compared to the longer isoform. PacBio long read sequencing did not identify the longest isoforms probably due to lower abundance. In contrast, *Tmx1* expressed longer isoform, through the increased use of a more distal site, even though the shorter isoform remained dominant (**Figure 2D**). While being correctly identified with Pacbio, this isoform was not annotated in the reference. The S100bp gene showed an increased use of its most distal site, the long 3’UTR isoform remaining dominant (**Figure S2**). Functional annotation (**Figure 2E**) showed, overall, the enrichment of APA transcripts in organelle, membrane trafficking pathways or intracellular transport suggesting a role for APA in cellular transcript localisation through possible regulation of their exposure to co-factors such as RNA-binding proteins or miRNAs. We further noticed the enrichment of transcripts with lengthened 3’ UTRs in the IL-1 pathway, which was observed only in the transition between 1 and 2 divisions. Finally, we also observed the shortening and lengthening of 3’ UTRs of transcripts associated with metabolism only between 1 and 2 divisions in activated samples supporting the notion that the metabolism keeps changing upon cell proliferation, but division is needed to initiate some 3’ UTR shortening/lengthening events.

**Figure 2.**
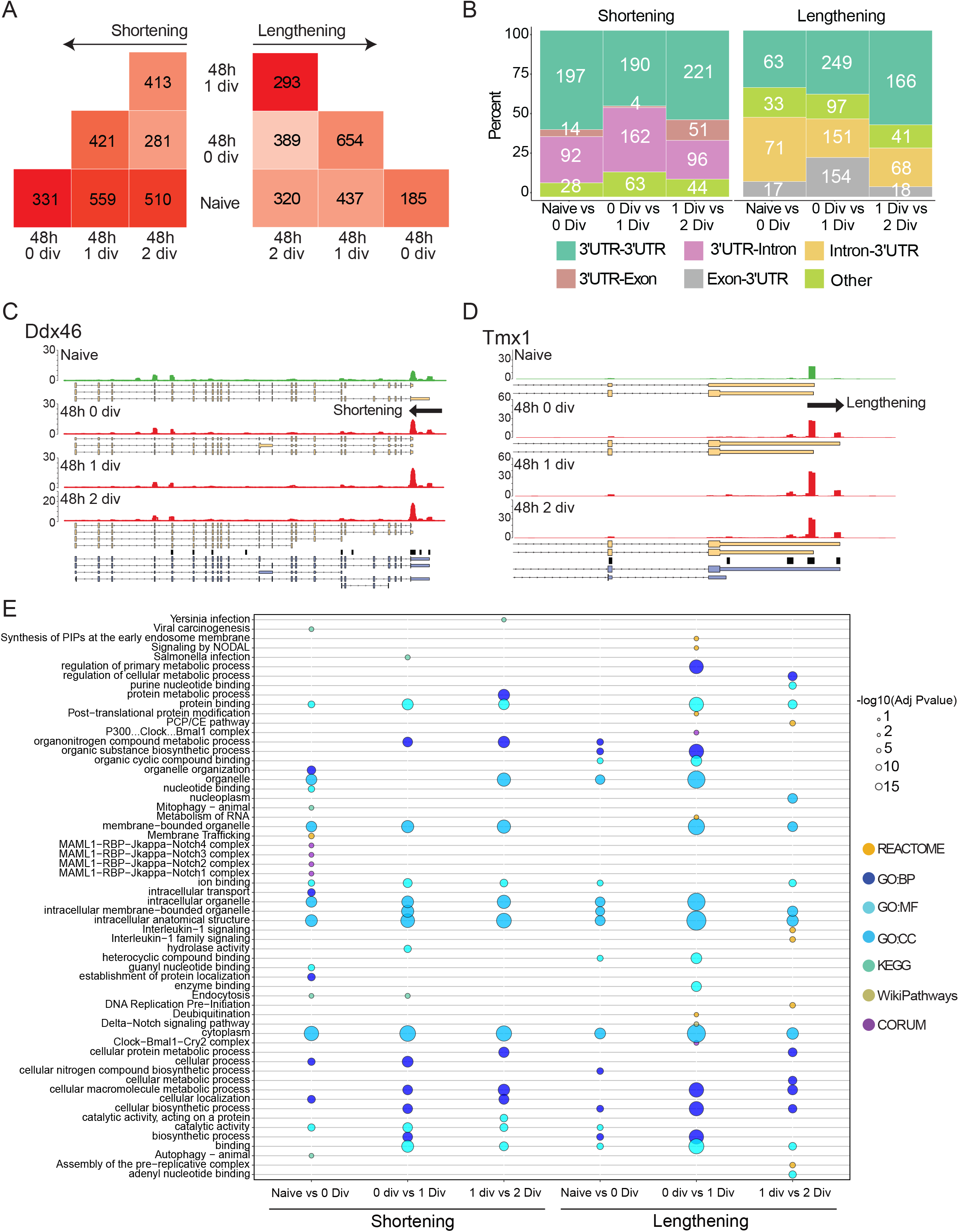
Identification of alternative polyA site usage. A) Number of shortening and lengthening events identified in each comparison. Dark color indicates higher numbers, light color indicates lower numbers. B) Classification of APA events per type of transition. C,D) Example of Ddx46 transcripts which are shortened and Tmx1 transcripts which are lengthened upon activation. Arrow indicates event direction. Yellow boxes represent isoforms identified using PacBio long read sequencing, blue boxes represent Gencode (v25) isoforms and blackboxes indicate PAS. E) Functional annotations of genes associated with an APA event. Dot colors indicate source types. Dot size represents -log10(adjusted Pvalues).

### Dynamic of alternative polyA site usage

While our and previous datasets demonstrate that global changes in 3’UTR length occur in T cells that are activated and dividing *in vitro* (Sandberg *et al*., 2008; Gruber *et al*., 2016), it is yet unknown in which T cell subpopulations these changes occur. When naïve T cells are activated they enter a phase of massive proliferation (clonal expansion). At the end of the immune response, the clonal population collapses (attrition), leaving a memory population of more quiescent T cells. We thus sought to determine how the polyadenylation landscape is remodelled as cells exit proliferation and become memory T cells. To this end, we sorted three different T cell populations from C57BL/6 mice: naïve T cells (CD4^+^ CD25^-^ CD62L^+^ CD44^-^), regulatory T cells (CD4^+^ CD25^+^), and effector memory T cells (CD4^+^ CD25^-^ CD62L^-^ CD44^+^) (**Figure S3A**). After RNA extraction, we generated a second batch of 3’ ends library by A-seq2 and compared them with the first batch (**Figure 3A**). The batch effect was limited, the variance within the four naïve samples was low (<8%) and therefore, we did not correct the data for batch effect and considered all naïve samples as replicates. PCA using RPM normalised read count over polyA sites indicated that CD4^+^ regulatory and memory T cells were more similar to naïve CD4^+^ T cells compared to 48h activated CD4^+^ T cells. Comparison of significant APA events number (FDR < 0.05) reflected the relationships observed in the PCA (**Figure 3B**): differences in polyA site usage were smaller between naïve and memory or regulatory T cells than between either of these and activated T cells. This was the case for both, shortening and lengthening, respectively. We annotated genes in which an APA event occurred between the second division following activation and acquisition of a memory and regulatory T cell phenotype (**Figure 3C**).

**Figure 3.**
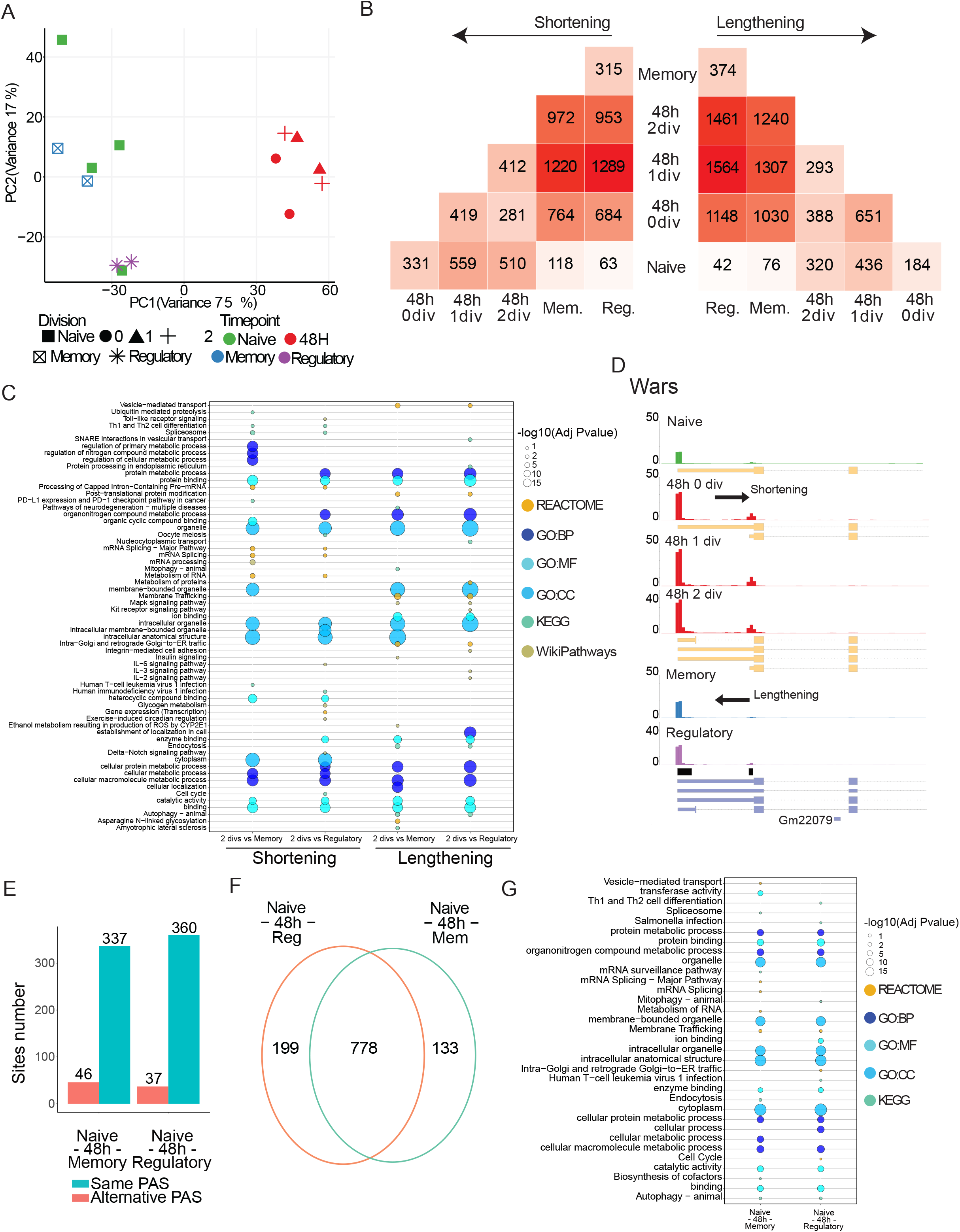
Dynamic of alternative polyA site usage. A) PCA resulting from A-seq2 read coverage over the final peak set using naïve, all 48h activated, memory and regulatory T cells. B) Number of shortening and lengthening events identified in each comparison. Dark color indicates higher numbers, light color indicates lower numbers. C) Functional annotations of genes associated to an APA event between activated 2 divisions and either memory or regulatory states. D) Example of 3’UTR Wars gene showing reverted length across activation and effector stages. E) Concordance of APA sites usage in genes associated with a reverting event. F) Venn diagram showing overlap of genes presenting a reverting pattern. G) Functional annotations of genes associated with a reverted APA event.

Organelle related pathways were enriched in both shortened and lengthened transcripts in these comparisons suggesting intense membrane trafficking changes as observed before (Grumont *et al*., 2004). Some pathways were specific to shortening events (for example, Th1 and Th2 cell differentiation, ubiquitin proteolysis or metabolism of RNA) whereas others were specific to lengthening events (for example, vesicle mediated transport, endocytosis, Processing of Capped Intron−Containing Pre−mRNA, Mapk signaling pathway). Compared to activated cells, regulatory T cells exhibited 3’ UTRs lengthening of transcripts enriched in the IL-2 and IL-3 pathways whereas shortening genes were more enriched for the IL-6 pathway. Thus, our data suggested that 3’ UTR modulation may differentially affect various T cell subsets, an observation in line with the notion that APA is context-dependent (Mitschka and Mayr, 2022).

To illustrate the dynamics of APA on a specific gene, we chose the tryptophanyl-tRNA synthetase (*Wars*), which was associated with a reversible remodelling of 3’ UTRs (**Figure 3D**). At the level of the entire transcriptome, the diversity of polyA sites used across all T cell populations was low: in 87% of the cases the dynamic of 3’ UTR shortening and lengthening involved the same APA sites (**Figure 3E**). Transcripts with reversible 3’UTR usage in either memory or regulatory T cell differentiation (**Figure S3B**) were very similar (778 transcripts of 1110) between these cell populations (**Figure 3F**). 199 and 133 transcripts underwent 3’UTR length changes specifically in one of the two cell populations (regulatory and memory CD4^+^ T cells). We functionally annotated those specific genes (**Figure 3G**). Similarly to memory T cells, regulatory T cells showed enrichment in organelle organisation. In addition, regulatory T cells functional annotations supported modification of gene expression and chromatin, suggesting the establishment of different regulatory programs in Tregs (Vignali, Collison and Workman, 2008). Altogether, these data showed that poly(A) site usage changes transiently during CD4^+^ T cells activation, proliferation and differentiation, and is associated with very specific functions during those phases.

### APA events and transcript abundance regulation

While many recent studies have focused on the mechanisms regulating the choice of poly(A) sites (Bucheli *et al*., 2007; Larochelle, Hunyadkürti and Bachand, 2017; Gruber *et al*., 2018; Grassi *et al*., 2019), decoupling transcriptional and post-transcriptional effects in the regulation of isoform abundance remains a key challenge. We interrogated our dataset by comparing it to a previously generated orthogonal total mRNA sequencing dataset generated from naïve and 48h activated CD4^+^ T cells (Dölz *et al*., 2022). While the activation period of this dataset matched with our study, it did not discriminate cells by division. To mimic this, we merged reads of all 48h activated A-seq2 samples and created 4 pseudo-replicates, accordingly with naïve samples (see METHODS). We first identified genes associated with an event by comparing naïve and all 48h activated samples (**Figure 4A**). We identified more shortening than lengthening events: 443 shortening events and 288 lengthening events. Similarly to the first analysis (**Figure 2B**), the majority of events involved sites located in 3’UTR or in intronic regions. Estimated levels of gene expression from A-seq2 and mRNAseq data were overall well correlated for naïve T cells and 48h after activation (Spearman r = 0.77 ; p < 2.2e−16 in naïve and r = 0.79 ; p < 2.2e−16 in activated samples) (**Figure 4B**). These results were in line with the observation that APA usage estimates from these two approaches correlate enough to be used by bulk RNAseq data based tools (Ha, Blencowe and Morris, 2018; Ye *et al*., 2018; Wang and Tian, 2020; Gerber, Schratt and Germain, 2021). To annotate transcriptional changes in genes that also exhibited APA, we performed a differential gene expression analysis and a differential transcript usage using the mRNAseq data (**Figure 4C**). 48h post activation, we observed an important rewiring of the transcriptome: 5,659 genes were differentially expressed (DEG) (False Discovery Rate (FDR) < 0.01; absolute logFold Change (FC) > 1 ; **Figure S4**). Of the 443 genes with shortened 3’ UTRs, 137 were also DEG (31%), 49 were associated with differential transcript usage (DTU) (11%) and 42 (9%) were both DEG and DTU. Of the 288 genes with lengthened 3’ UTRs, 85 were also DEG (30%), 30 were associated with DTU (10%) and 28 were both DEG and DTU (9%).

**Figure 4.**
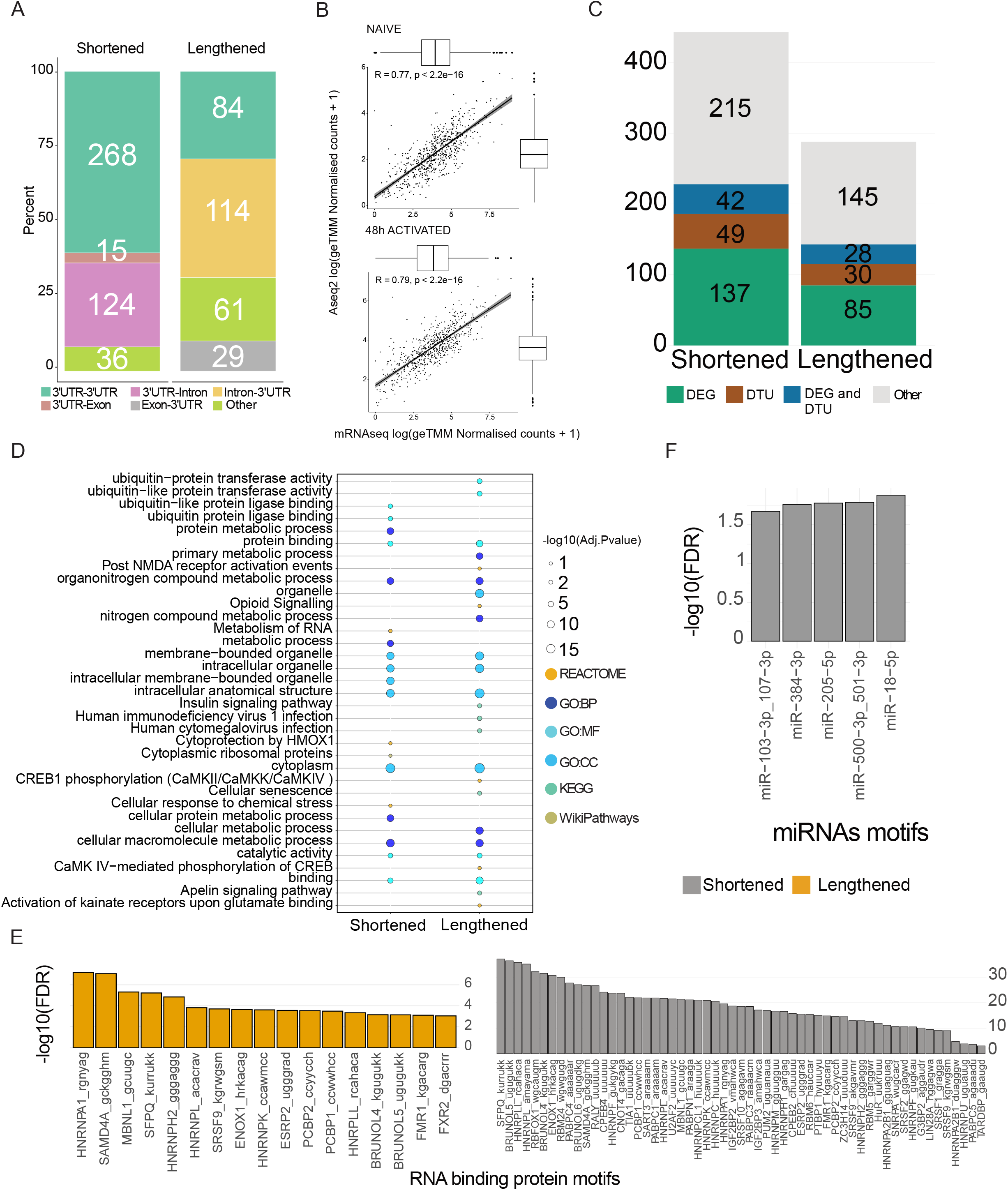
APA events and transcript abundance regulation. A) Classification of APA events per type of transition. B) Pearson correlation of mRNAseq (Dolz et al.) and Aseq2 gene coverage at naive (top) and 48h after activation (bottom). Reported values are log transformed Normalised counts + 1. Boxplots indicate value distribution and their mean. C) Barplot reporting number of APA genes being also differentially expressed (DEG), having a differential transcript usage (DTU) both or none. D) Functional annotations of genes associated with an APA event when comparing naive and pooled 48h activated samples. E) Barplot showing enrichment for RBPs within alternative 3’UTR by type of event, shortening (grey) or lengthening (yellow). Colour legend indicates -log10 (FDR). F) Barplot showing enrichment for miRNA binding sites within alternative 3’UTR by type of event, shortening (grey) or lengthening (yellow). Colour legend indicates -log 10 (p value).

Functional annotation of APA genes when comparing naive and all combined activated samples (not taking into account the cell division status) (**Figure 4D**) revealed biological pathways related to organelle biology shared both by APA genes being shortened or lengthened. To note, ubiquitin−protein transferase activity pathway was enriched only in lengthened transcripts whereas ubiquitin−protein ligase activity pathway only in shortened transcripts. The adequate abundance and localisation of transcripts and proteins, crucial for the cell, is regulated at different levels. Longer transcript isoforms can be more exposed to post-transcriptional regulation because they can contain more cis-regulatory miRNA (Sandberg *et al*., 2008) or RNA binding protein recognition sites (Jiang and Coller, 2012; Gruber and Zavolan, 2019). To analyse whether shortening or lengthening events remove or add putative RBP sites, we scanned dynamic 3’UTR regions for RNA motifs known to be recognized by RBPs, all available in RBPmap database (Paz *et al*., 2014). Out of the 92 tested motifs, 84 were significantly enriched, in the alternatively included region of either shortened or lengthened 3’UTRs, compared to the 3’UTR regions of genes that did not exhibit APA in T cells (Fisher’s exact test corrected for multi-testing; FDR < 0.001) (**Figure 4E**). Among the top significant associations, HNRPLL is already known to promote memory T cell RNA rearrangements (Wu *et al*., 2008), and in the control of alternative splicing in T cells (Gaudreau *et al*., 2012), ZC3H14 has a known role in controlling the length of polyA tails (Kelly *et al*., 2014). We repeated this analysis with a set of well characterised and conserved miRNA binding sites available in the Mouse TargetScan database (McGeary *et al*., 2019). 5/218 miRNA binding sites were significantly enriched relative to background (Fisher’s exact test; p value < 0.05; background was genes expressed in CD4+ T cells) in shortened genes only (**Figure 4F**). Among the miRNAs expressed in T cells, we found enrichment for miR-18-5p which is part of the miR-17-92 cluster and known to be involved in T cell activation and differentiation (Dölz *et al*., 2022) and miR-103-3p/107-3p and miR-500-3p/501-3p which are not characterised in T cells yet. Present analysis suggested a well-balanced system in which APA usage has limited effect on global gene expression.

## Discussion/Conclusion

In this study, we have analysed the dynamics of 3’ UTR length during T cell activation and differentiation. While we previously assessed alternative polyadenylation in pan-T cells after 72 hours of *in vitro* stimulation (Gruber *et al*., 2014), here we sought to investigate the 3’UTR landscape in purified naïve CD4^+^ T cells and to compare it to 48h *in vitro* activated CD4^+^ T cells as well as ex vivo purified effector memory and Treg cells. In addition, we aimed to resolve effects on APA induced by T cell activation versus proliferation. To this end we used a division-dependent resolution and we combined direct detection of 3’ ends using A-Seq2 with PacBio long-read sequencing. This allowed us to better define the 3’UTR dynamics, associated with particular post-transcriptional regulators. We observed that, on average, there were slightly more 3’UTR shortening events than 3’UTR lengthening, in line with previous work (Hsin *et al*., 2018), but different from what was observed upon sustained T cell activation (Sandberg *et al*., 2008). Our analysis of effector memory T cells and Treg cells suggests that APA is transient since the 3’ UTR site usage of these cells was more similar to naïve T cells than activated T cells. While the activation and subsequent proliferation of naïve CD4^+^ T cells resulted in global changes in the 3’UTR landscape, we were interested in the steady state situation of distinct T cell populations. By looking at the 3’UTR landscapes of naïve and memory T cells we were comparing a polyclonal pre- to a post-expansion population. One limitation of this study comes from the 0 division activated cells which were phenotypically heterogeneous. It might therefore be that the number of APA genes would increase with stronger activation. Thus, our data does not allow a thorough conclusion to differentiate the effect of T cell activation vs cell cycle entry. Nevertheless, by pairwise comparison of cell division stages, we could show an unprecedented APA dynamic usage at the cell division level. Our data also suggest that effector/memory and regulatory CD4+ T cells shared polyA sites usage over naïve CD4 T cells and point to a yet unrecognised reversibility of global 3’UTR dynamic. Alternative polyadenylation therefore exhibits a form of plasticity that is inducible upon T cell activation and might be reversible in vivo after differentiation into memory cells. Comparisons with published data using a mRNA dataset showed that APA genes were also differentially expressed between naïve and activated samples, and around 10% presented a differential transcript usage, demonstrating a marginal effect or association of alternative poly-adenylation with global gene expression and transcript usage. Importantly, APA does not equally affect all transcripts (Gruber *et al*., 2016). It has been shown that miRNAs and RBPs coevolved with APA events (Karginov, Ménoret and Vella, 2022), resulting in an intricate balance of regulation versus counter-regulation to control cell fate decisions. The availability of 3’UTR binding sites on specific transcripts and the differential expression of miRNAs and RBPs participate in the competition between different regulators of mRNA abundance supporting a regulatory network that might serve as a post-transcriptional determinant of cellular function. Future work on transcriptional regulation in different T cell types and differentiation stages, but also B cell populations, is needed to reveal the involvement of alternative polyadenylation within tightly regulated differentiation steps and receptor activation thresholds controlled in all ranks to ensure a balanced and working immune system. Technology-wise, approaches such as nascent-RNA sequencing coupled with 3’ end sequencing at the single cell level will help to answer unsolved questions.

## Acknowledgments and funding

The authors would like to thank Robert Ivanek (DBM Bioinformatics Core Facility) and all lab members of the Jeker and Zavolan lab for critical discussions, Georges Martin for help with the ASeq2 protocol and Anna Devaux for text contribution and reviewing, the DBM Flow Cytometry Facility for high quality cell sorting, the DBM Animal Facility team for in-house breeding and excellent animal care and the D-BSSE genomics facility for generating and sequencing RNA libraries. This project has received funding from the Swiss National Science Foundation (SNSF Professorship PP00P3_144860 to LTJ), the National Institute of Allergy and Infectious Diseases of the National Institutes of Health, USA, under Award Number R56/R01AI106923 and the European Research Council (ERC) under the European Union’s Horizon 2020 research and innovation programme (grant agreement No. 818806).

## Conflict of interest

LTJ is a co-founder and board member of and holds equity in Cimeio Therapeutics AG a biotech company developing engineered cellular therapies. LTJ’s activities related to Cimeio are unrelated to this study. All other authors have no CoI to declare.

## Author contributions

Conceptualization: O.G., LJ; Methodology: D.S., O.G., R.S., M.Z.; Samples collection & data generation: O.G., R.M. ; Software and algorithm implementation: D.S., O.G., R.S., M.Z.; Data curation: D.S., O.G., R.S., M.Z.; Data analysis, D.S., O.G., R.S., M.Z.; Investigation, D.S., O.G., R.S., M.Z.; Writing groups – Original Draft: D.S., O.G., R.S., M.Z.; Writing – Review & Editing, everyone; Figures preparation: D.S., O.G.; Funding Acquisition, L.J. and M.Z. ; Supervision, L.J. and M.Z., Project administration, L.J. and M.Z.

## Methods

### Data and code availability

The A-seq2 datasets generated in this study are available under accession ID GSE183424. PacBio long-read sequencing data are accessible under accession ID GSE209604. mRNAseq dataset (Dölz *et al*., 2022) are available under accession ID GSE140568 and miRNAseq dataset will be available in the future under accession ID GSE203633. Code is available at https://gitlab.com/JekerLab/dynamic-3p-utr-changes-in-cd4-t-cell.

### T cell isolation, stimulation and sorting

Spleens and lymph nodes were isolated from male C57BL/6N mice at an age of 6-10 weeks. Animal work was done in accordance with the federal and cantonal laws of Switzerland. All organs were collected on ice in complete T cell medium (RPMI 1640 (Sigma), 10% heat-inactivated FCS (Atlanta Biologicals), 2mM Glutamax (Gibco), 50μM β-mercaptoethanol (Gibco), 10mM HEPES (Sigma) and non-essential amino acids (Gibco)) and organs were mashed with a syringe plunger through 45μm filters in medium under sterile conditions. Cell suspensions were transferred to 15 ml tubes and centrifuged for 5 min at 400 g, 4°C. Red blood cell lysis was performed with ACK buffer for 5 min at RT and stopped by addition of medium. Naïve CD4^+^ T cells were isolated with EasySep Mouse Naïve CD4^+^ T Cell Isolation Kit (STEMCELL Technologies Inc) according to manufacturer’s recommendations. For naïve T cell activation 12 well plates (Corning) were coated with 1ml monoclonal anti-CD3 (2μg/ml, 2C11, BioXcell) in 500 μl PBS for 2 hours at 37°C, 5% CO_2_. Purified naïve CD4^+^ T cells were labelled with 2μl CellTracerViolet dye (CTV; in DMSO) in 2ml PBS for 20 min at 37°C and labelling reaction was stopped by addition of 10 ml complete medium. Cells were plated in a density of 4 × 10^6^ CTV-labelled naïve T cells per coated 12 well or 6 well plates in 2 ml or 4 ml, respectively, plus anti-CD28 antibodies (1μg/ml, PV-1, BioXcell) and incubated at 37°C with 5% CO_2_ for 48 hours. 4-6 × 10^6^ purified naïve CD4^+^ T cells were resuspended in Trizol on the day of T cell purification (day 0). After 48 hours of cell culture cells were harvested and stained with 1μg/ml propidium iodine briefly before sorting. Cells were sorted by their distinct proliferation peaks (peak 0, peak 1 and peak 2) on FACS-Aria and Influx Cell Sorters (BD Biosciences). Cells were kept on ice and after washing with cold PBS were immediately resuspended in Trizol for RNA extraction. One portion of the cells were used for surface phenotype staining with the fluorochrome-conjugated mAbs: anti-CD4 (clone RM4-5), anti-CD25 (PC61), CD62L (MEL-14), CD44 (IM7), CD69 (H1.2F3), CD16/32 (93, all Biolegend). Cells were measured on a LSR-Fortessa (BD Biosciences) and data was analysed using FlowJo (Treestar). For isolation of steady-state populations of CD4+ naïve, memory and regulatory T cells, cells were pre-isolated with EasyStep Mouse CD4^+^ T cell isolation Kit (STEMCELL Technologies Inc) and stained with the fluorochrome-conjugated mAbs targeting CD4, CD44, CD62L and CD25 before being sorted on FACS-Aria and Influx Cell Sorters (BD Biosciences). Purified populations were washed with cold PBS and immediately resuspended in Trizol for subsequent RNA extraction.

### RNA Extraction

Isolated cells were quickly processed and washed with cold PBS before resuspension in 1ml Trizol. Samples were left at room temperature for 5 min and frozen at -20°C. For RNA extraction, Trizol samples were thawed on ice and 100 μl 1-Bromo-3-Chloropropane was added before mixing and incubation for 15 min at RT. Samples were centrifuged for 15 min at 12000g, 4°C and the aqueous phase was transferred to a fresh tube before mixing with 1 volume isopropanol and a subsequent incubation of 10 min at RT. Samples were then centrifuged for 15 min at 12000g, 4°C and the RNA pellet was washed with 1 ml ethanol followed by a centrifugation for 30 min at 12000g, 4°C. The supernatant was discarded and the pellets briefly dried before the RNA dissolved in dH2O and stored at -80°C.

### A-seq2 protocol

Briefly, poly(A)+ RNAs were isolated from RNA samples and fragmented by alkaline hydrolysis. Next, adaptors were ligated to the 5’ end of enriched poly(A)+ RNAs and reverse transcription was carried out with a bio-dU-dT(25) RT-primer. After USER-enzyme digestion and purification, a second adaptor was ligated to 5’ ends of the resulting cDNA. Both adaptors were finally used to PCR-amplify libraries with barcoded primers and used for RNA sequencing (Gruber *et al*., 2016; Martin *et al*., 2017).

### Preprocessing of A-seq2 libraries

We first perform quality analysis and read trimming of A-seq2 read libraries with fastp (v.0.19.7) and the following parameters: minimum length required (-l) = 15, UMIs were integrated by enabling -U, placed in the read 1 (--umi_loc)=read1 and formed of first 7 bases of each reads (--umi_len=7) and finally max N number limit was set to 2 (--n_base_limit). Mapping of the trimmed reads on mm10 genome was achieved with STAR (Dobin *et al*., 2013) (v.2.7.3) by asking end to end alignment (--alignEndsType EndToEnd) and no multi mapped reads (--outFilterMultimapNmax 1). Finally, we used UMIs to remove duplicated reads with umi_tools (v.1.0.0)(Smith, Heger and Sudbery, 2017) *dedup* (--umi-separator=“:” --extract-umi-method=“read_id” --method=“directional”)

### Poly(A) site quantification and events identification and functional annotations

#### Poly(A) site quantification and genomic annotation

Read enriched regions (“peaks”) were called using Fseq (v.1.84) (Boyle *et al*., 2008) with fragment size (-f) of 50, feature length (-l) of 150 and threshold (-t) of 6. Bins around peaks with less than 10 reads were removed to refine peaks. Within each condition, peaks from each replicates were intersected using BEDTools (v.2.27.1) (Quinlan and Hall, 2010) and resulting intersected peak sets were combined into a master set of peaks using DiffBind R package (v.2.16.2)(Ross-Innes *et al*., 2012) and a minimum overlap of 1 (peaks need to be present in at least one intersected peakset). Master set of peaks was then sloped by 200bp both sides and overlapped with Poly(A) sites from the PolyAsite resource for mouse version r2.0 (https://www.polyasite.unibas.ch/atlas) (Herrmann *et al*., 2019). We filtered out peaks with a size greater than 2000bp and smaller than 10bp. A-seq2 reads were quantified using featureCounts function from Rsubread package (v.2.4.3) (Liao, Smyth and Shi, 2019). Peaks were categorised based on their genomic locations (5’UTR, 3’UTR, Exon, Intron, Intergenic) using Gencode VM25 annotations (https://www.gencodegenes.org/mouse/release_M25.html).

### Event identification and filtering

To identify events (shortening or lengthening), we used PolyA-miner (https://github.com/LiuzLab/PolyA-miner) (Yalamanchili *et al*., 2020). Parameters were set as follows: polyA annotations file (-pa) contained the sites identified above, reference fasta sequence (-fasta) was downloaded from Ensembl (http://ftp.ensembl.org/pub/release-94/fasta/mus_musculus/dna/Mus_musculus.GRCm38.dna.toplevel.fa.gz) and reference genes bed file (-bed) was created from Ensembl gff3 file (v94). The types of events were stringently defined to fulfil two conditions: transcript is shortened if sample B proximal projection was greater than sample A proximal projection and if sample B distal projection was smaller than sample A distal projection. The opposite conditions identified as a lengthened gene. Events were then filtered on FDR (5%). For each pairwise comparison, MaxAPASwitch position from PolyA-miner output was retained as the most used position on condition B whereas the most covered position was considered as the most used APA site in condition A. This was done according to type of event and strand. The DNA sequence between sites from condition A and B was further used for miRNA and RBP binding/motif analysis.

### Functional annotation of APA genes

Functional annotation was carried out using the gprofiler2 R package (v.0.2.0)(Raudvere *et al*., 2019), model organism was set to “mmusculus”, “fdr” correction method and the following sources were interrogated: GO:BP, GO:MF, GO:CC, KEGG, REAC, MIRNA, CORUM and WP. A 5% FDR threshold was applied and the top 5 annotations within each of the sources were plotted.

### mRNAseq processing and analysis

mRNAseq data count tables were downloaded from GSE140568 and processed similarly to Dolz et al. Briefly, here are the main steps. The GTF file used along the processing steps was extracted from Gencode VM25.

### mRNA acquisition and sequencing

2.5*10^5 cells were washed with PBS, resuspended in 200μl TRI Reagent and RNA was extracted from Trizol-samples with a Zymo Direct-zol kit which includes DNAse treatment. RNA quality was assessed with a Fragment Analyzer (Advanced Analytical) and RNA-seq library preparation was performed using Illumina Truseq stranded kit. Sequencing was performed on an Illumina NextSeq 500 machine to produce single-end 76-mers reads. All steps were performed at the Genomics Facility Basel (ETH Zurich).

### Gene-level quantification

Read quality was assessed with the FastQC tool (v.0.11.5)(http://www.bioinformatics.babraham.ac.uk/projects/fastqc). Reads were mapped to the mouse genome (UCSC version mm10) with STAR (v.2.5.2a)(Dobin *et al*., 2013) with default parameters, except filtering out reads mapping to more than 10 genomic locations (outFilterMultimapNmax=10), reporting only one hit in the final alignment for multi mappers (outSAMmultNmax=1) and filtering reads without evidence in the spliced junction table (outFilterType-“BySJout”).

### Differential analysis

Read alignment quality was evaluated using the qQCReport function of the R Bioconductor package QuasR (v.1.18) (Gaidatzis *et al*., 2015). Gene expression was quantified using the qCount function of QuasR as the number of reads (5’ends) overlapping with the exons of each gene assuming an exon union model (using the UCSC knownGenes annotation downloaded on 2015-12-18). The R Bioconductor package edgeR (v.3.28) (Robinson, McCarthy and Smyth, 2010) was used for differential gene expression analysis. Between-samples normalisation was done using the TMM method. Only genes with CPM (counts per million mapped reads) values more than 1 in at least 4 samples (the number of biological replicates) were retained. A generalised linear model including a genotype effect, an activation effect, and a replicate effect (nested within genotype) was fitted to the raw counts (function glmFit), and differential expression was tested using likelihood ratio tests (function glmLRT). P-values were adjusted by controlling the false-discovery rate (Benjamini-Hochberg method) and genes with a FDR lower than 1% were considered differentially expressed.

### Differential transcript usage (DTU)

We used the DRIMSeq R Bioconductor package (v.1.20.0)(Nowicka and Robinson, 2016) to perform DTU on mRNAseq data. First, for each 4 replicates, in naïve and 48h activated samples, salmon (v.1.5.0) quantification was performed at transcript level using *quant* function and the following parameters: -1 A --validateMappings --seqBias --gcBias --posBias --softclip --numBootstraps 100. A full model comparing naïve and activated samples was applied, post-hoc filtering was executed (it improves the FDR and overall FDR control by setting the p-values and adjusted p-values for transcripts with small per-sample proportion SD to 1). Finally a stage-wise procedure (stageR R bioconductor package v.1.14.0) was used to adjust P-values.

### miRNA quantification

miRNAseq data were mapped to the mm10 genome using Bowtie2 and --very-sensitive-local parameter. UMIs were counted using a custom python script.

### miRNAs and RBPs binding sites analysis

Background regions were 3’UTR regions of genes considered as expressed in T cells (log2(CPM+1) > 0 in at least naïve or 48h activated).

### miRNAs binding site enrichment analysis

Genome coordinates of Predicted Conserved Targets were retrieved from Target Scan Mouse database (Release 7.2) (Agarwal *et al*., 2015). This BED file contains genome (mm10) locations of mouse predicted (conserved) targets of conserved miRNA families and associated score (context++ score percentile). Enrichment for each individual miRNA family against shortening and lengthening regions from each comparison was computed and Fisher Exact Test p-value was reported.

### RBPs motif enrichment analysis

DNA sequences of dynamic 3’UTR regions were extracted and converted into RNA sequences. They were scanned with RBPmap (v.1.2) (Paz *et al*., 2014) RBP motifs, converted into Transfac format. Motif enrichment analysis was performed with Analyse Motif Enrichment (*ame*) from MEME suite (v.5.3.3) (McLeay and Bailey, 2010). Parameters were as follows: --scoring avg --method fisher --hit-lo-fraction 0.25 --evalue-report-threshold 10.0 --control --shuffle-- --kmer 2. FDR threshold of 0.1% was applied.

### PacBio Long read sequencing

Three SMRTbell libraries (Naïve, 48h activated 0 division and 2 divisions) from Trizol isolated high-quality RNA samples (>500 ng total RNA) were sequenced on a Sequel SMRT Cells 1M for a general survey of full-length isoforms in a transcriptome with moderate to high expression levels. SMRTbell libraries and sequencing were achieved by the Department of Biosystems Science and Engineering -ETH (D-BSSE). Tools from PacBio & Bioconda were used to process these data (https://github.com/PacificBiosciences/pbbioconda). Consensus sequences (.ccs) were generated from your raw subread data by using ccs (--min-rq 0.9). Full-length reads were generated after primer removal and demultiplexing using lima --ccs. Then we removed polyA tails and artificial concatemers with isoseq3 refine (--require-polya --min-polya-length 20) and clustered consensus sequences to generate transcriptome fasta with isoseq3 cluster (--use-qvs). High-Quality Full-Length Polished Isoforms were mapped to the genome and collapsed into transcripts based on genomic mapping using pbmm2 align (--preset ISOSEQ). We then used isoseq3 collapse to generate fastq and gff files. Transcripts were annotated using SQANTI3 (v0.1) (Tardaguila *et al*., 2018) (--aligner_choice minimap2) and providing list of mouse polyA motifs retrieved from PolyASite database (https://polyasite.unibas.ch/atlas), transcript abundance from isoseq3 output and CAGE peaks from Fantom5 (https://fantom.gsc.riken.jp/5/datafiles/reprocessed/mm10_latest/extra/CAGE_peaks/).

## Notes

### Competing Interest Statement

The authors have declared no competing interest.

