## Supplemental figure 1 for "T helper cells exhibit a dynamic and reversible 3’UTR landscape"

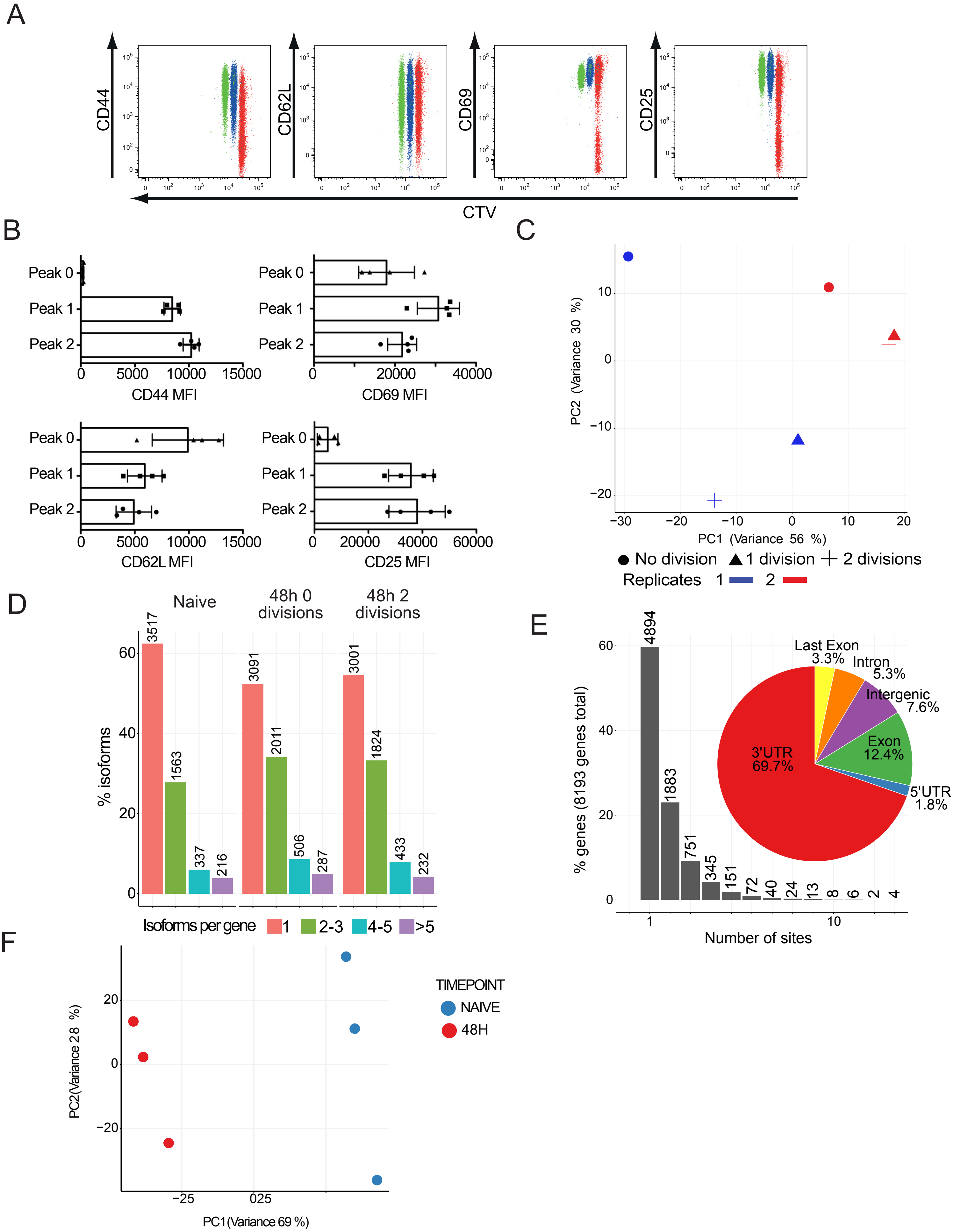


Supplemental Figure 1 related to figure 1. A) Representative immunphenotyping for activation markers CD44, CD62L, CD69, and CD25 separated by proliferation peak populations (red = no division (peak 0), blue = 1 division (peak 1), green 2 divisions (peak2)) after 48 hours in vitro stimulation. B) Mean fluorescence intensities (MFIs) are shown for the respective markers expressed by the given proliferation peak population analyzed by flow cytometry in four independent samples (mean +/- SD). C) PCA resulting from read coverage over the final peak set showing activated samples only and the different cell divisions. D) Distribution of isoform per genes identified with long read sequencing PacBio data. E-F) Summary of Hsin data reprocessing. E) Barplot representing distribution of APA sites per genes. Pie plot shows genomic localization distribution of APA sites. F) PCA resulting from read coverage over the final peak set identified with Hsin et al. dataset.
