## Supplemental figure 2 for "T helper cells exhibit a dynamic and reversible 3’UTR landscape"

# S100bp

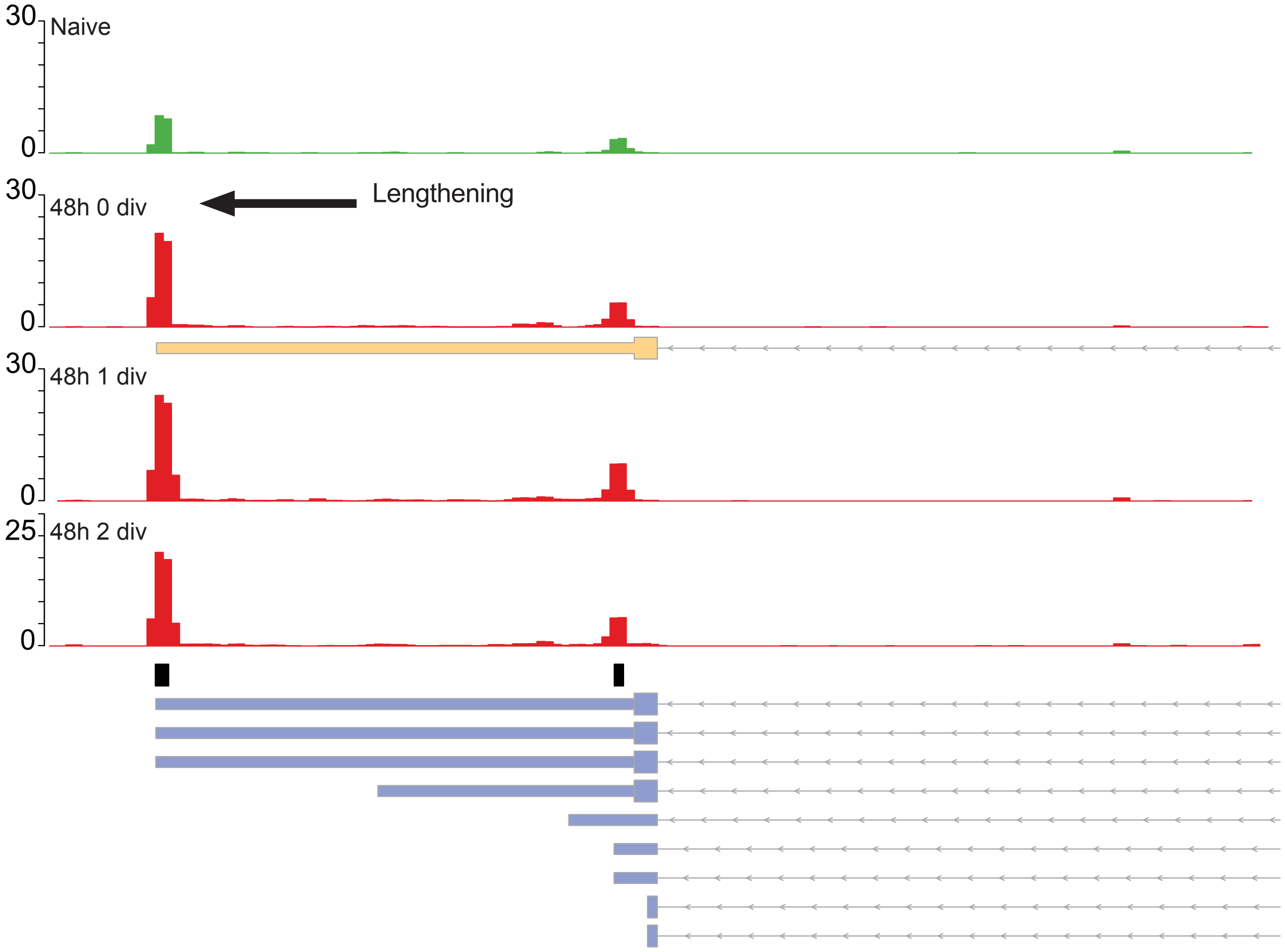


Supplemental Figure 2 related to figure 2. Example of S100bp transcript which is lengthened upon activation. Arrow indicates event direction. Yellow boxes represent isoform identified using PacBio long read sequencing, blue boxes represent Gencode (v25) isoforms and black boxes indicate PAS.
