## Supplemental figure 3 for "T helper cells exhibit a dynamic and reversible 3’UTR landscape"

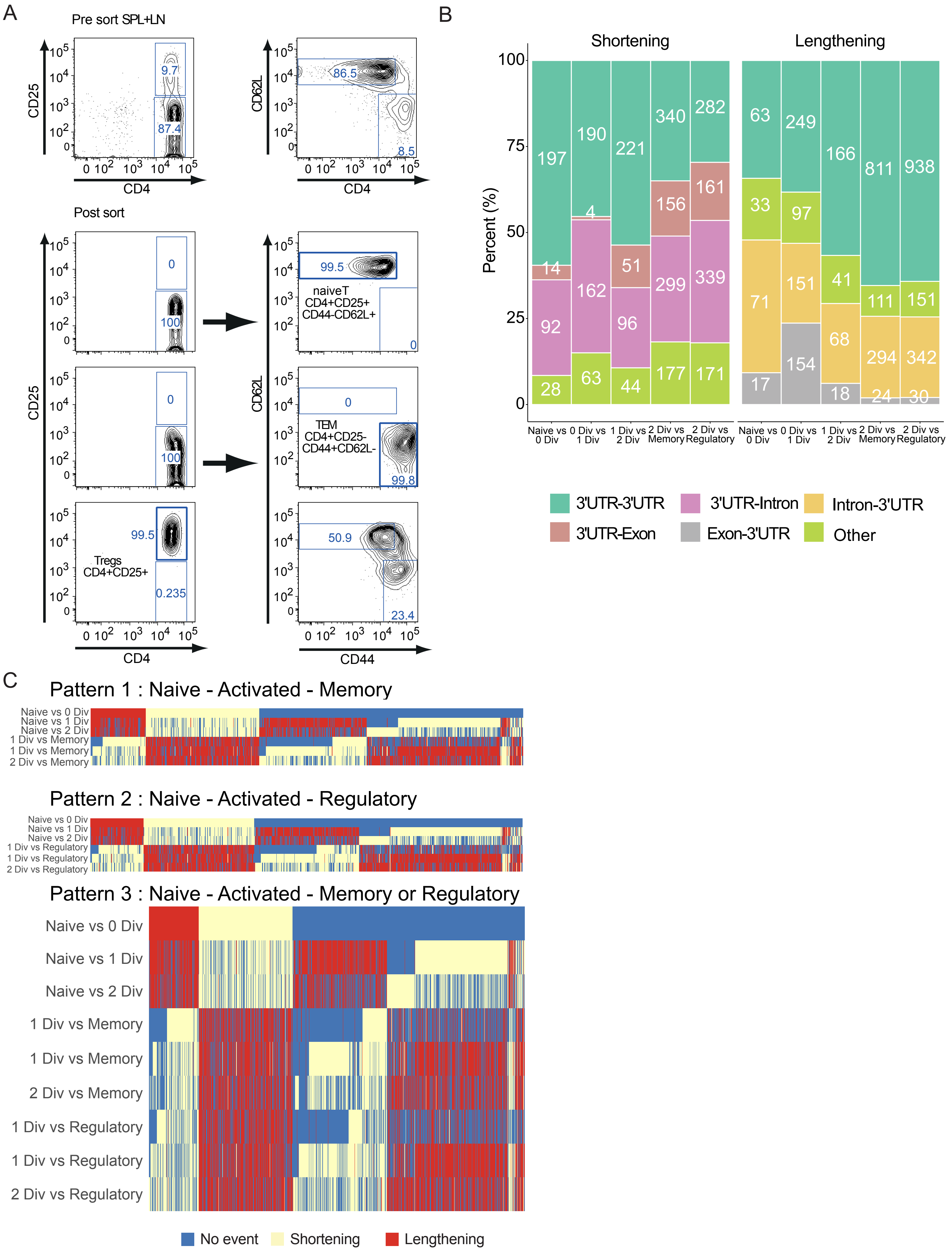


Supplemental Figure 3 related to figure 3. A) Representative immune-phenotyping of CD4+ T cells for activation markers CD44, CD62L, CD69, CD25 after 48 hours of in vitro stimulation. B) Classification of APA events per type of transition. C) Overview of all genes having a reverted pattern, e.g. being shortened/lengthened between naïve and one of the 48h activated populations, and then being lengthened/shortened in either memory or regulatory
