## Supplemental figure 4 for "T helper cells exhibit a dynamic and reversible 3’UTR landscape"

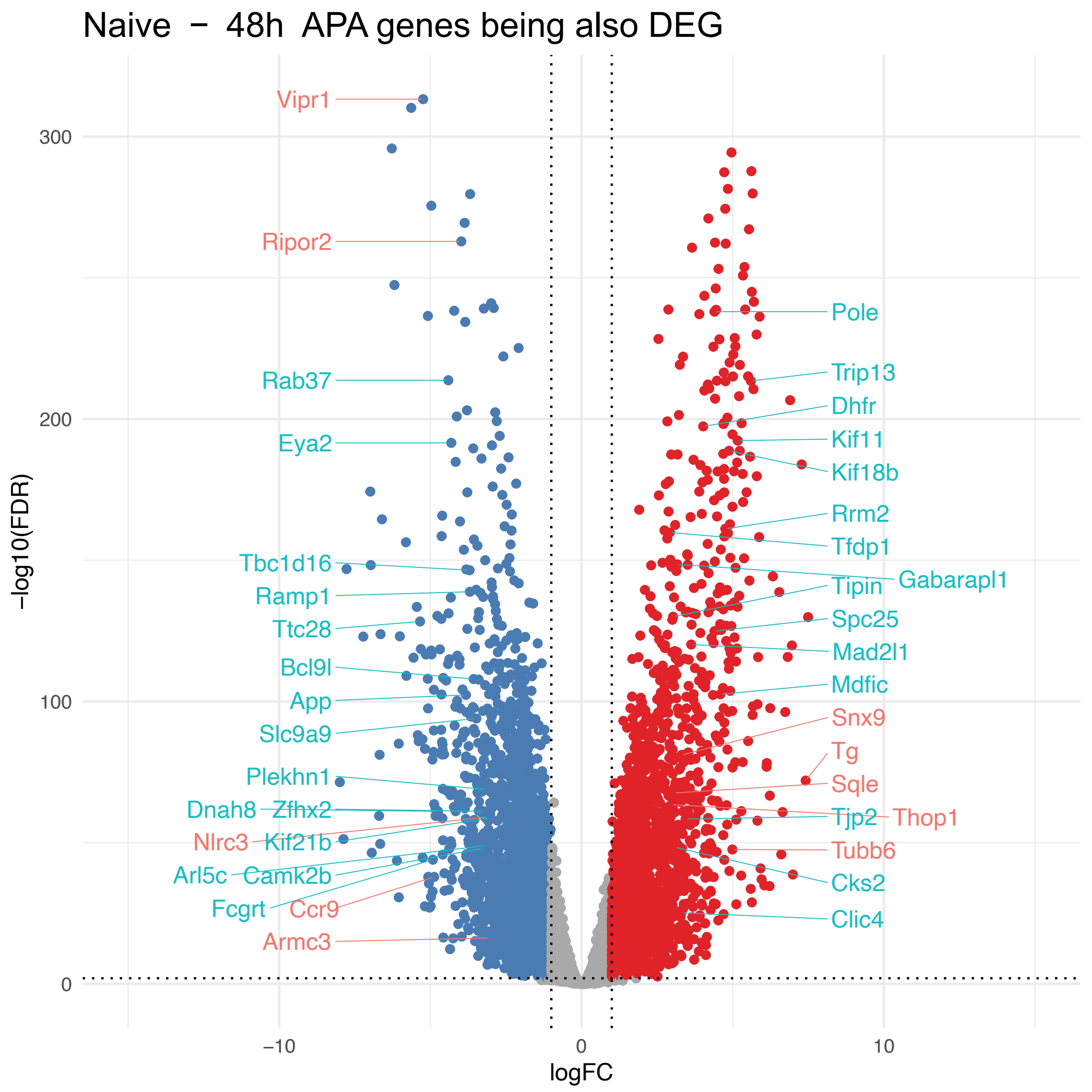


Supplemental Figure 4 related to figure 4. Volcano plot resulting from mRNA differential analysis (Dolz et al.). Blue and red dots indicate genes with false discovery rate (FDR) < 0.01 and absolute log fold change (FC) > 1. APA genes being also DEG are indicated: ligth blue indicates shortening event and red lengthening event.
