## Supplemental table 1 for "T helper cells exhibit a dynamic and reversible 3’UTR landscape"

| | | Naive | 0 division | 2 divisions |
|---|---|---|---|---|
| Unique | Isoforms | **10,527** | **12,602** | **11,109** |
| | Genes | **5,644** | **5,902** | **5,499** |
| Gene classification | Annotated | 4,751 | 5,582 | 5,162 |
| | Novel | 893 | 320 | 337 |
| Characterization of transcripts based on splice junctions | FSM (Full Splice Match) | 5,552 | 7,324 | 6,550 |
| | ISM (Incomplete Splice Match) | 1,112 | 1,978 | 1,878 |
| | NIC (Novel In Catalog) | 1,824 | 1,761 | 1,407 |
| | NNC (Novel Not in Catalog) | 921 | 1,047 | 793 |
| | Genic / Genomic | 98 | 73 | 55 |
| | Antisense | 343 | 156 | 156 |
| | Fusion | 15 | 8 | 6 |
| | Intergenic | 662 | 253 | 264 |
| | Genic Intron | 0 | 2 | 0 |
| Splice junction classification | Known canonical | 44,857 (97.47%) | 53,239 (97.6%) | 48,206 (98.05%) |
| | Known non-canonical | 37 (0.08%) | 29 (0.05%) | 32 (0.07%) |
| | Novel canonical | 879 (1.91%) | 1,020 (1.87%) | 709 (1.44%) |
| | Novel non-canonical | 247 (0.54%) | 261 (0.48%) | 217 (0.44%) |
